# CRISP: Correlation-Refined Image Segmentation Process

**DOI:** 10.1101/2024.08.23.609461

**Authors:** Jennifer K. Briggs, Erli Jin, Matthew J. Merrins, Richard K.P. Benninger

## Abstract

Calcium imaging enables real-time recording of cellular activity across various biological contexts. To assess the activity of individual cells, scientists typically manually outline the cells based on visual inspection. This manual cell masking introduces potential user error. To ameliorate this error, we developed the Correlation-Refined Image Segmentation Process (CRISP), a two-part automated algorithm designed to both enhance the accuracy of user-drawn cell masks and to automatically identify cell masks. We developed and tested CRISP on calcium images of densely packed β-cells within the islet of Langerhans. Because these β-cells are densely packed within the islet, traditional clustering-based image segmentation methods struggle to identify individual cell outlines. Using β-cells isolated from two different mouse phenotypes and imaged on two different confocal microscopes, we show that CRISP is generalizable and accurate. To test the benefit of using CRISP in functional biological analyses, we show that CRISP improves accuracy of functional network analysis and utilize CRISP to characterize the distribution of β-cell size.

## Introduction

Calcium imaging is widely used for quantifying cellular activity across many different fields of biology^1^. A benefit of calcium imaging is the ability to capture the activity of multiple cells at the same time. In neuroscience, calcium imaging has been used to elucidate neuron communication, response to stimulus, and neuronal function^1^. Alternatively, in the pancreatic islet, which is central to diabetes, is composed of thousands of β-cells that communicate electrically to secrete insulin in a coordinated, pulsatile fashion. Therefore, measuring the activity and coordination of many β-cells at one time has become a rich topic to study^2–4^.

To study the activity and coordination of cells using calcium imaging, cell masks must be created to distinguish individual cells. The image intensity of all pixels within a mask is then averaged to create the calcium time course for each cell. In sparse cell populations, such as some neurons, automated cell masking methods based on intensity thresholds can be used^5,6^. However, these methods do not typically perform well in organoids with densely packed cells, such as the pancreatic islet or cardiomyocytes. To study these systems, a researcher takes a series of time-stacked calcium images and manually outlines each cell of interest to create a mask. This methodology presents two possible problems. First, when assessing the coordination between multiple cells, it is essential that the pixels within each cell outline are within the cell of interest. If the researcher inadvertently includes pixels within the cell mask that were truly a part of the neighboring cell, the coordination between the two cells would be over-reported. Alternatively, in practice, researchers typically outline cell masks from either the first calcium image or the average of all images in time. Given the large number of pixels within each cell mask and the large number of cell masks generated by the researcher, small outline mistakes are common even for expert and careful researchers. The second methodical problem is that manually circling cells in a thorough enough manner to avoid including neighboring pixels is fatiguing for the researcher. This is especially the case if there are many cells to manually outline, researcher fatigue causes less accurate outlines, making the error in the cell mask heteroskedastic (more error as time and fatigue increases), and possible confounding conclusions.

In this manuscript, we develop two automated algorithms based on a single principle to ameliorate these methodological problems arising from using manually identified cell masks for time series analysis.

## Results

### Pixels within a cell are highly correlated compared to those outside of the cell

We first tested our hypothesis that pixels within a cell were more highly correlated with each other than with pixels outside the cell, even for a highly coupled system such as β-cells within the islet of Langerhans (**Fig 1a)**. We therefore conducted calcium imaging of GCamP6-labelled β-cells within an islet using a Nikon A1R confocal microscope (**Fig 1b)**. We then created a circular mask with a radius of 30 pixels centered at a β-cell nucleus and extracted the calcium time course **(Fig 1b)**. Strikingly, a colormap showing the correlation coefficient between the center pixels and all other pixels can be used to distinguish a single cell, indicating that pixels within a cell are more correlated with each other than those outside of the cell (**Fig 1c)**. Using this finding, we developed CRISP – Correlation Refined Imaged Segmentation Process. CRISP can be used for two tasks. Automated cell mask refinement (**Fig 1c, right top)** uses CRISP to refine manually drawn cell masks. Automated semi-minor axis identification (**Fig 1c, right bottom)** uses CRISP to identify in an unbiased manner the largest circle that contains only pixels within a cell.

**Figure 1:**
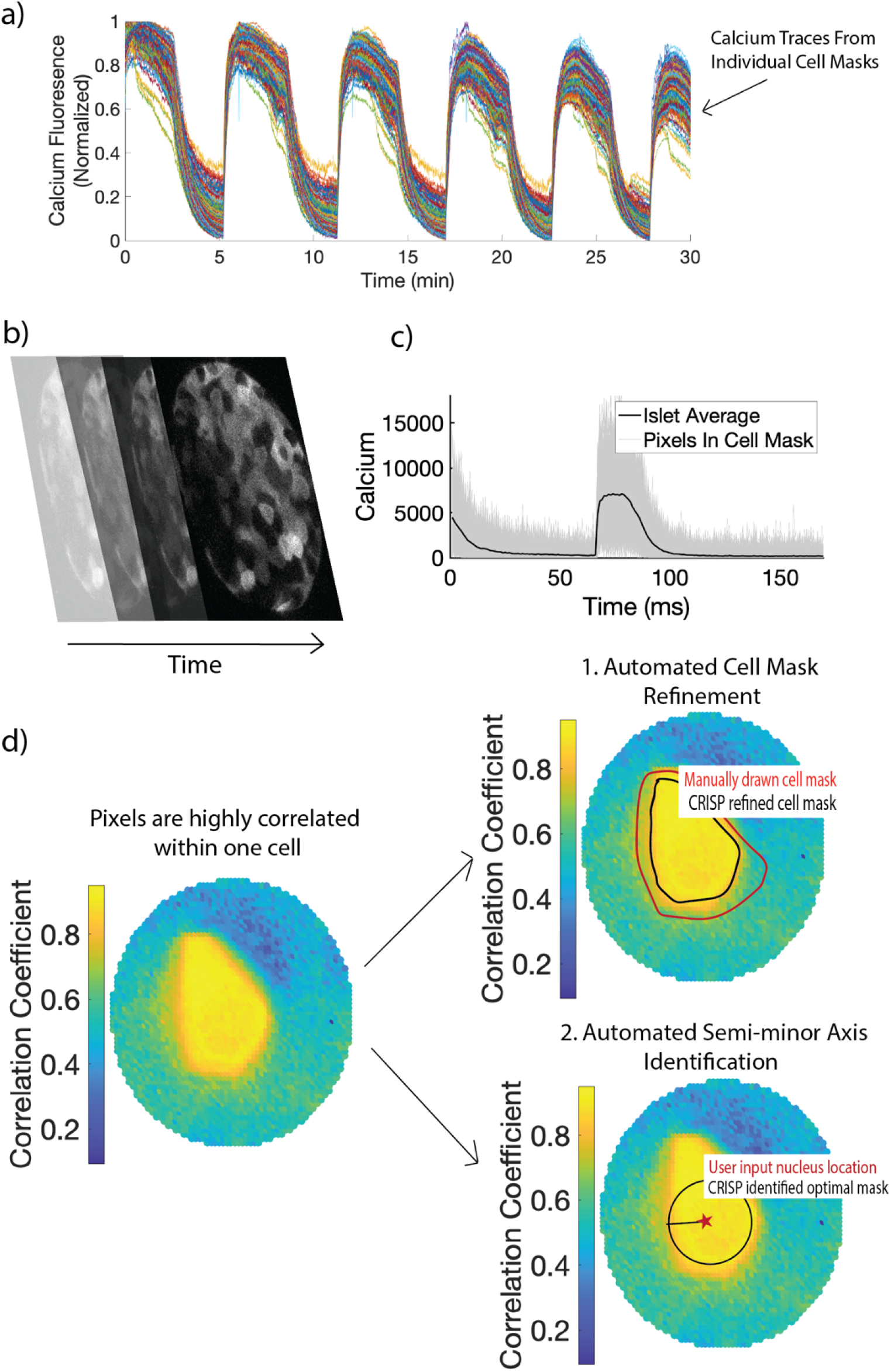
(a) Example time courses from β-cell masks. Each line indicates calcium fluorescence from a single β-cell mask. (b) Example of calcium image stacks of pancreatic islet. (c) Resultant calcium time course of all pixels within a manually outlined cell mask (d) Left: Color plot showing the correlation between the nucleus and the rest of the pixels in the region of interest. Cell outline can be visualized only based on the correlation color plot. Using this observation, two algorithms are created. Right: 1. Automated mask refinement takes user defined mask (drawn in red) and removes uncorrelated pixels that are not a part of the cell to create the CRISP refined cell mask (black). 2. Automated semi-minor axis identification begins with the user input nucleus location (red star) and identifies the largest circle radius that captures pixels only within the cell of interest.

### Automatic cell mask refinement with CRISP

GoodTo assess CRISP accuracy and identify the optimal correlation threshold for removing pixels, “bad”, “medium”, and “good” masks were manually circled with differing levels of precision (see methods) for 45 cells (15 cells for three islets) **(Fig 2a)**. CRISP was used to refine medium and bad masks and performance was assessed by comparing CRISP-masks to good mask. Visual inspection for a bad mask shows that CRISP correctly identifies mostly correct pixels to remove from the manually outlined mask (**Fig 2b)**. CRISP mask refinement resulted in an area under the ROC of 0.84±0.03 for bad masks and 0.81±0.05 **(Fig 2c)**, indicating that CRISP is effective at mask refinement. An optimal correlation threshold of 0.25 was chosen to maximize true positive rate (correctly keeping pixels that belong to the cell), while minimizing false positive rate (incorrectly keeping pixels that belong to the cell). Before CRISP, there was high variance in the time courses between pixels within cell masks **(Fig 2d)**. CRISP successfully removed these pixels with high variance (or low correlation), thereby decreasing the noise in individual cell masks **(Fig 2e)**.

**Figure 2:**
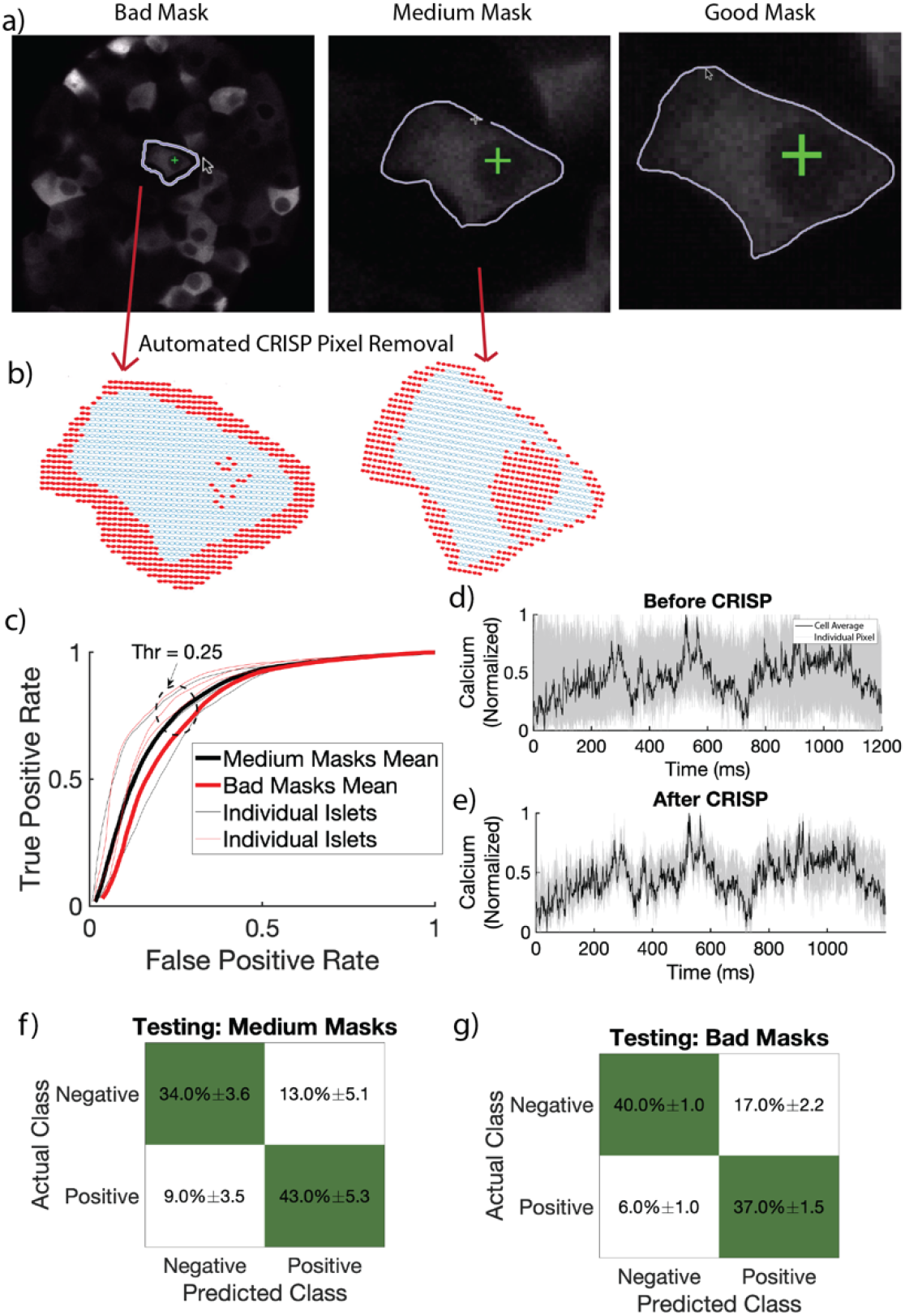
Automated Cell Mask Refinement. a) Representative bad (circled without zooming), medium (circled with zoom), and good (circled carefully with zoom) masks. b) CRISP removed pixels (red) from representative bad and medium masks in (a). c) Receiver operating characteristic curve (ROC) of four islets with fifteen β-cells circled per islet. Red lines indicate ROC curves for bad user defined masks. Black lines indicate ROC curves for medium user defined masks. Dark lines indicate average over four islets, while light lines indicate individual islets. Light lines indicate individual islets. Average area under the curve **0.83.** d) Calcium time course of all pixels in a bad mask before CRISP. Grey lines indicate individual pixels. Black line indicates average calcium over the full cell. e) As in c but after CRISP. f) Confusion matrix showing performance of testing data set for medium masks. g) As in e for bad masks.

To ensure generalizability, we tested CRISP using a separate data set, originally reported in^7^. This data set was generated from a different microscope (LSM5Live) with lower resolution and microscope settings and a separate mouse genotype (Connexin 36 knockout mice). We tested CRISP on three islets, with 10 masked each (n = 30). For both medium (**Fig 2f)** and bad masks **(Fig 2g)**, CRISP correctly estimated 77% of pixels. Considering the total number of pixels characterized by CRISP was ∼1000 and that the testing set came from a completely different mouse model and microscope, this indicates that CRISP generalizes well to different systems and methods of calcium image data collection.

### CRISP cell refinement improves correlation coefficient accuracy

Functional network analysis is commonly used in fields of islet biology and neuroscience to understand how synchronized different cells are to one another. CRISP automatic cell refinement was run on medium and bad masks for 60 cells from four islets **(Fig 3a)**. To test if CRISP could improve the accuracy of network analysis results, we calculated the correlation coefficient between each cell within the islet **(Fig 3b)**. Medium masks resulted in an average correlation coefficient error (compared to good masks) of 0.0052. CRISP significantly decreased this error to 0.005 **(Fig 3c)**. Bad masks resulted in an average correlation coefficient error of 0.0067. This error resulted in an average difference of 1.3 connections, with a max of 6 connections difference. CRISP significantly decreased this correlation error to 0.0057 **(Fig 3d)**. Therefore, CRISP can improve the accuracy of functional network theory calculations, particularly when the initial masks are poor.

**Figure 3:**
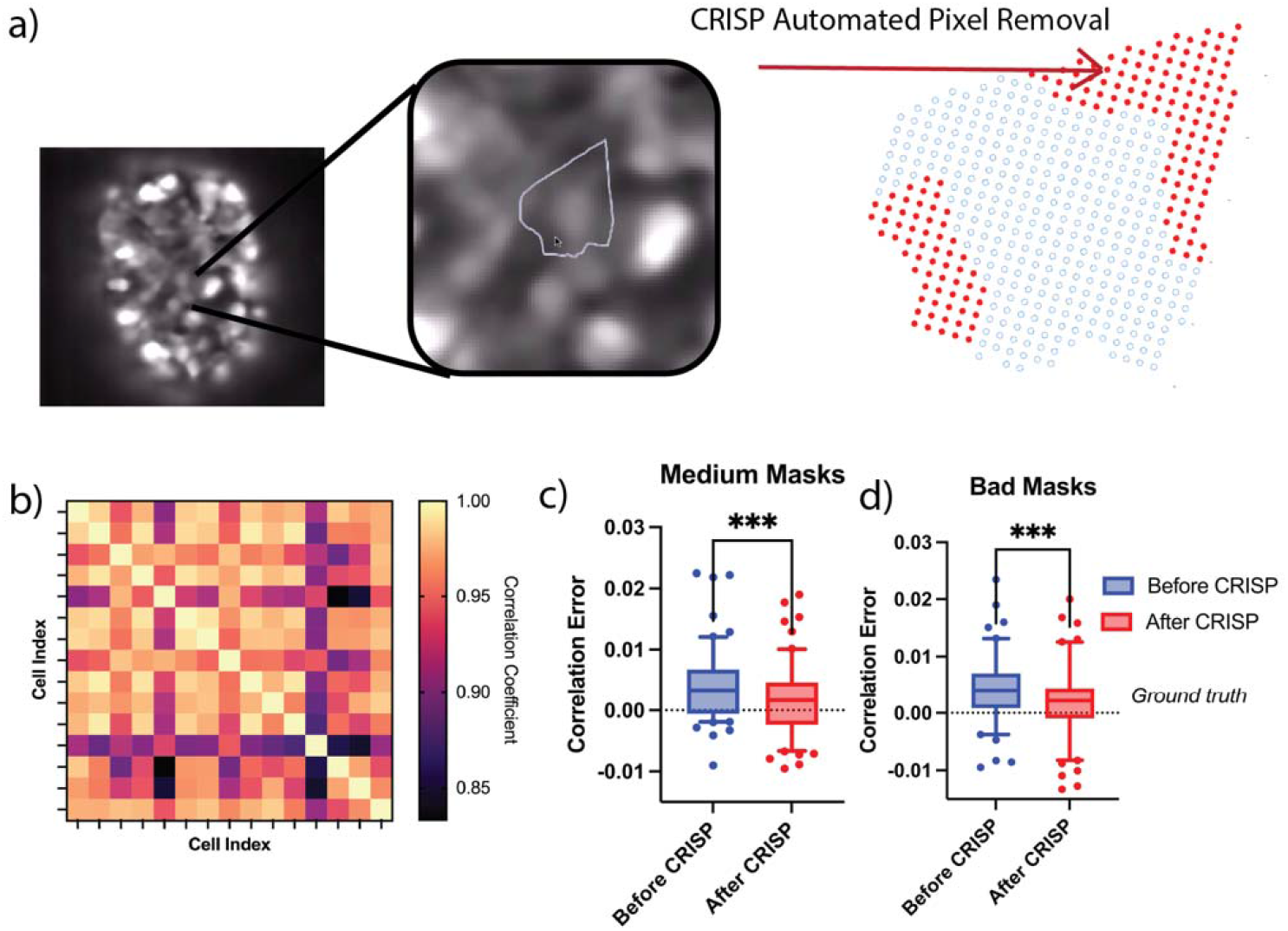
a) Example figure of islet with bad user defined mask. Red pixels (right) are pixels correctly removed from mask by CRISP. b) Heat map showing correlation coefficient between 15 cells in an islet. c) Difference between correlation coefficient from medium and good masks before CRISP (blue) and after CRISP (red) p = 0.001, d) As in h but for bad masks p < 0.001.

### Automatic semi-minor axis identification with CRISP

In high throughput settings or three-dimensional settings, it is not realistic to manually outline every cell. Instead, automated calcium extraction methods (such as for Bitplane Imaris Software) require the user to input the radius of the cell mask. To determine the largest radius of a circular cell mask centered at the nucleus that contained only pixels from within the cell, we developed a two-step algorithm. We manually drew masks around 10 cells from 5 islets to serve as training data. In the first algorithmic step, we identified the optimal correlation coefficient that maximized the correctly labeled cells within a large radius (40 pixels).

We first calculated the optimal correlation threshold (**R**_**th**_, **Algorithm 2**). For each cell in the training set, we created a circular annulus (or field of view) with a radius of 40 pixels around the cell. This radius was to ensure that the full cell would be well within the annulus. We set all pixels within a 2-pixel radius from the center of the cell as the baseline to compare to. This baseline is defined differently than for automated cell mask refinement because the approximate cell mask size is not known. We then calculated the correlation coefficient between every cell in the annulus and the baseline pixels over many fixed thresholds. We use a fixed correlation threshold (as opposed to the standard deviation-based threshold used in automatic cell refinement) because the large annulus likely contains pixels from other cells, and thus a standard-deviation based threshold may be biased towards other cells in the annulus. The optimal correlation threshold **R**_**th**_, was 0.818, with an area under the receiver operating curve of 0.8 **(Fig 4a)**. Upon visual inspection in some annuli, the cell was accurately identified with only the correlation threshold **(Fig 4b)**. However, in other annuli, large numbers of pixels were incorrectly labeled to be inside of the cell **(Fig 4c)**.

**Figure 4:**
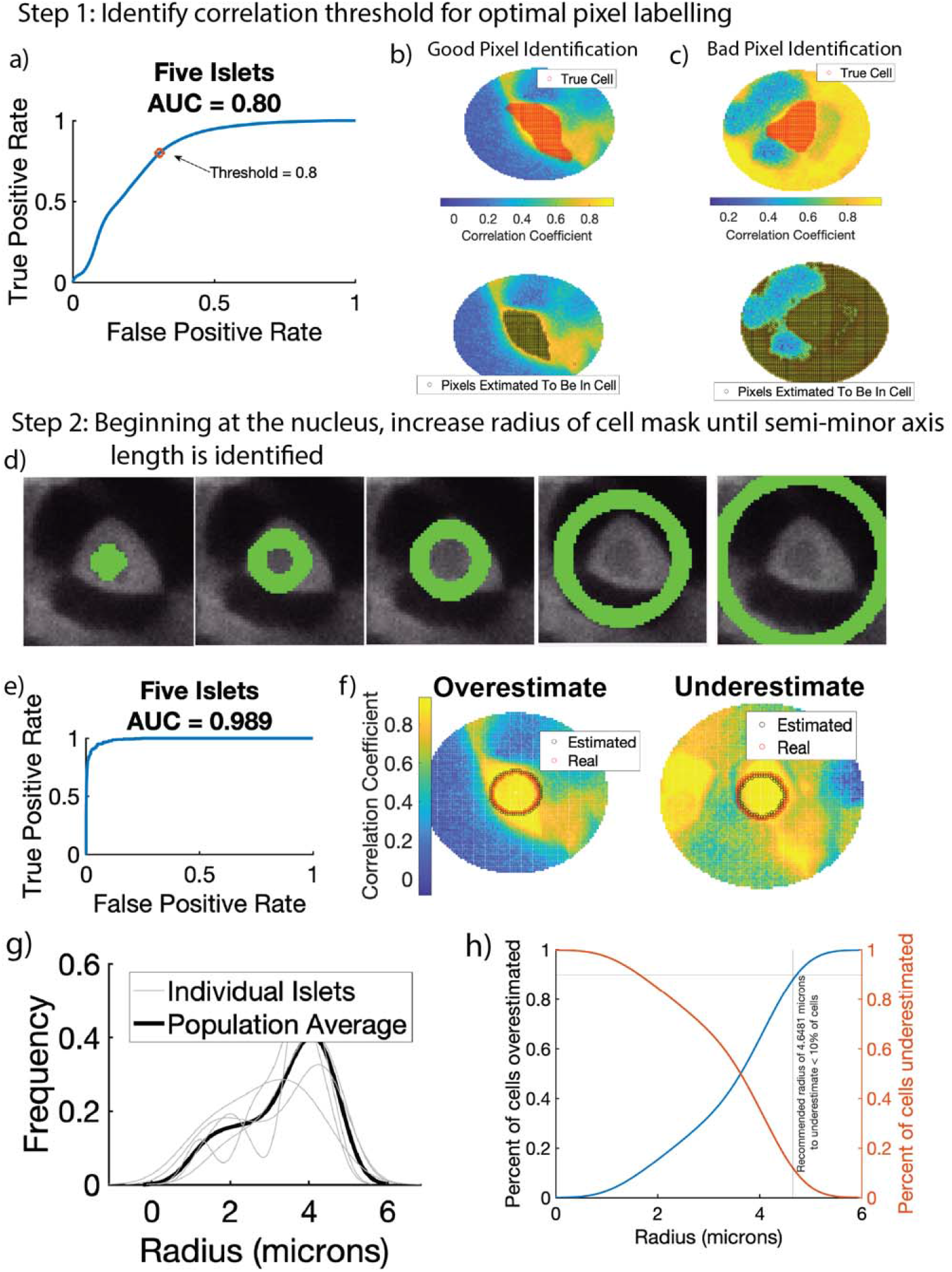
Automated Semi-minor axis identification in two steps. In step 1, the correlation between every pixel in a large region and the user-defined nucleus is calculated. Receiver operating characteristic curve (ROC) had an area under the curve of 0.8. Threshold of 0.8 was identified for step 2. b) Representative example of bad pixel identification. Top shows correct cell mask (red) overlaid on a correlation color map. Bottom shows pixels estimated to be inside the cell (black). c) Representative example of good pixel identification. Correct cell mask (red, top) is closely estimated by CRISP (black, bottom). In step 2, after threshold of 0.8 is identified in step 1, CRISP begins with a small annulus surrounding the nucleus and calculates a score (see methods). CRISP then iteratively increases annulus radius until score threshold is reached. d) Left shows initial anulus radius, middle shows annulus with where CRISP identified optimal radius, right shows annulus that are too large and minimize score. e) ROC curve for CRISP semi-minor axis identification algorithm. Area under the curve is 0.989. f) Example of annulus that is underestimated by CRISP. g) Example of annulus that is overestimated by CRISP. Using CRSIP, the semi-minor axis length was estimated for five islets with approximately 150 cells. a) Distribution of radius of β-cells. b) Recommendation for high throughput β-cell calcium analysis where a single radius is defined for all β-cells.

We then created the iterative annulus expansion step. In this step, the annulus was initialized at 2 pixels **(Fig 4d, left)**, the score function for that annulus was computed and the annulus radius was increased. To identify the optimal score threshold (**p**_**+**_), we calculated the score for all training cells up to an annulus radius of 40 pixels **(Fig 4d, right)**. The area under the ROC curve of 0.989 indicates that CRISP is highly accurate at semi-minor axis identification **(Fig 4e)**. CRISP did over- and underestimated multiple cell semi-minor axes, but only by one or two pixels (**Fig 4f)**.

We used CRISP to assess the average semi-minor axis length (radius) of 150 β-cells from five islets. Each of the five islets resulted in a roughly similar radius distribution, which was positively skewed with an average radius of 4.0866 microns (**Fig 4g)**. Only 30% of cells were larger than 4.0866 microns. For very high throughput analyses, such as 3D light-sheet imaging, running CRISP for each cell may be unnecessarily computationally expensive. Instead, it would be optimal to identify a set radius that consistently captures the maximum possible number of pixels. After estimating the average radius, we identified an optimal radius of 4.65 microns, which underestimates approximately 10% of cells **(Fig 4h)**.

## Discussion

Time course imaging is used throughout fields of biology. The increase in microscope resolution has provided the opportunity to study activity from individual cells. When studying this activity a cell mask must be created. In cases where cells are densely packed, such as β-cells in the islet, this mask must be created manually. This act of manual masking induces user error and can be time-consuming. We sought to remove this potential error by developing an algorithm: Correlation-Refined Image Segmentation Process (CRISP) that utilizes intracell correlations to refine cell masks and automatically circle cells.

CRISP cell mask refinement algorithm resulted in what is typically considered an “excellent” ROC area under the curve^8^. When testing CRISP on an independent dataset, the average correct labeling was approximately 77%. Considering that the labels were on individual pixels, and the resultant calcium trace is an average of all pixels, 23% of incorrectly labeled pixels will have a negligible impact on the resultant calcium trace. Further, our testing set was calcium time courses taken between 2007-2008 with very different imaging technology than our training set (which was collected in 2023) and a different mouse model and different microscope modality (Zeiss line scanning vs Nikon confocal systems respectively). This was intentional to fully test the generalizability of CRISP. With these considerations in mind, 77% accuracy was particularly impressive.

Methods exist for fully automating cell masking using clustering algorithms^6^. These algorithms rely on differences in intensity for clustering and work well in systems where cells are sparsely packed (e.g., neurons). However, they tend to not perform well in systems where cells are densely clustered, such as islets or cardiomyocytes. Correlation-based methods also exist^9,10^. However, these methods utilize an iterative approach where individual pixels are added if under some correlation threshold, similar to Step 1 of Automated Semi-minor axis identification. We showed that these algorithms also do not work well in dense images because multiple highly correlated cells will be lumped together **(Fig 4c)**. By including an iterative annulus growth approach, we greatly increase the algorithm accuracy (AUC = 0.989).

This CRISP algorithm can be used to accomplish two tasks: automated cell mask refinement and automated semi-minor axis cell masking. The first task could be beneficial to remove errors induced by fatigue or multiple scientists masking a data set. We do not recommend that CRISP cell mask refinement be used as a replacement for careful and thoughtful cell masking, but rather an additional step to improve mask accuracy. The second task would be beneficial for times when the data are extremely plentiful and perfect masks are not necessary. In this case, CRISP can be used for rapid identification of the majority of pixels within each cell, removing the need for user-drawn masks.

## Methods

### Ca^2+^ Imaging

Islets were isolated from *Ins1-Cre:ROSA26*^*GCaMP6s/H2B-mCherry*^ mice as in^11^. The following day, islets were loaded in a QE1 quick exchange chamber (Warner Instruments) mounted on a Nikon A1R confocal microscope equipped with a 60×/1.42 NA oil objective. The imaging solution contained, in mM: 135 NaCl, 4.8 KCl, 2.5 CaCl_2_, 1.2 MgCl_2_, 20 HEPES, 10 mMol glucose. Excitation of GCaMP6s (488 nm, 20% power) and H2B-mCherry (561 nm, 10% power) was used with a 405/488/561 multi-band dichroic and emission filters (GCaMP6s: 525/50; H2-B-mCherry, 595/50) to report simultaneously report cytosolic Ca^2+^ levels and nuclear location. Timelapse imaging was performed at 0.125 Hz for 20 minutes.

### Testing set, first presented in^7^

Islet isolation from a Connexin 36 knockout mouse was performed following the methods of Scharp et al. and Stefan et al.,^12^ and the islets were maintained in RPMI medium with 10% fetal bovine serum and 11 mM glucose at 37°C in 5% CO2 for 24-48 hours before imaging. Isolated islets were stained with 4 mM Fluo-4 AM in imaging medium (125 mM NaCl, 5.7 mM KCl, 2.5 mM CaCl2, 1.2 mM MgCl2, 10 mM HEPES, 2 mM glucose, 0.1% BSA, pH 7.4) at room temperature for 1-3 hours. Imaging was conducted using a PDMS microfluidic device that stabilizes the islet and allows for rapid reagent changes. Fluorescence imaging was performed 15 minutes after increasing glucose from 2 mM to 11 mM using an LSM5Live microscope with a 20x 0.8 NA Fluar Objective and a 488 nm diode laser. The device was maintained at 37°C in a humidified, temperature-controlled chamber, with images acquired at 4-6 frames per second and average power minimized to 200 mW/cm^2^. A total of five islets were imaged. CRISP: Automated Cell Mask Refinement algorithm omitted two of these islets because of too much movement. CRISP: Semi-minor axis identification was robust to movement and utilized all five islets.

### Cell Masking

To test if CRISP improved the accuracy of manually drawn cell masks, manual masks were drawn around cells in three ways. Good masks were drawn by zooming in closely on the cell of interest and carefully outlining the exact cell. Each good mask took approximately 15 seconds to draw.

These masks were used to define the “true cell”. Medium masks were drawn after zooming in to the cell of interest but drawing around the cell without being careful. Each medium mask took approximately 5 seconds to draw. Bad masks were drawn without zooming into the cell. Each bad mask took approximately 1 second to draw.

### CRISP: Automated Cell Mask Refinement (Algorithm 1)

The CRISP algorithm compares the correlation coefficient between pixels with a baseline time series. This baseline time series is defined as the 10% of the pixels nearest to the center of the cell mask. The CRISP score for each pixel was defined as the correlation coefficient between the pixel and the baseline time series. To ensure generalizability across systems with different within-cell correlation (*r*), pixels were removed if they had a CRISP score less than the average CRISP score minus the standard deviation of all CRISP scores times a threshold (*R*_*th*_).

To identify the optimal threshold, bad and medium masks were run through CRISP using correlation thresholds between -2 and 0.3. Positive labels were defined as pixels that were labeled to be in the cell mask. For example, true positives were defined as pixels in the “good” cell mask and correctly labeled by CRISP to stay in the cell mask, whereas false positives were defined as pixels that were considered to be in the cell mask (e.g,. not removed) by CRISP but were not present in the “good” cell mask.

### CRISP: Semi-minor Axis Identification (Algorithm 2)

The baseline time series was defined as all cells within a circular annulus within a 2-pixel radius of the nucleus. The annulus was given a score function (eq. 1) based on the percent of pixels in the annulus (*p*_+_) with a correlation coefficient less than a threshold (R_th_) minus the annulus radius (*ϕ*) divided by 100. This penalty function ensures that the radius does not get too large in case the cell of interest is directly located next to another highly synchronized cell.

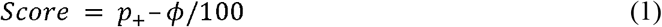

To identify the optimal score, the CRISP algorithm was run for thresholds (***S*_*th)*** between 0.7 and 0.95.

## Data availability

All calculations and image masking were conducted in MATLAB. All code is publicly available at: https://github.com/jenniferkbriggs/CRISP.

## Author Contributions

E.J. and R.K.P.B constructed the microscope and performed islet imaging experiments. J.K.B. developed algorithm and conducted analysis. J.K.B drafted the paper. All authors edited the paper. R.K.P.B and M.J.M provided resources

## Acknowledgements

We thank Barak Blum at the University of Wisconsin-Madison for providing Rosa26H2B-mCherry mice and the University of Wisconsin Optical Imaging Core for use of the spinning disk confocal. Jennifer K Briggs gratefully acknowledges support from NSF GRFP (DGE-1938058_Briggs). The Benninger laboratory gratefully acknowledges support from the NIH/NIDDK (R01DK106412, R01DK102950, R01DK140904 to R.K.P.B.) and the University of Colorado Diabetes Research center (P30 DK116073). The Merrins laboratory gratefully acknowledges support from the NIH/NIDDK (R01DK113103 and R01DK127637to M.J.M., and R01DK106412 to R.K.P.B.) and the United States Department of Veterans Affairs Biomedical Laboratory Research and Development Service (I01BX005113 to M.J.M.).

## Algorithms

### Algorithm 1

*Algorithm for CRISP automatic cell refinement*

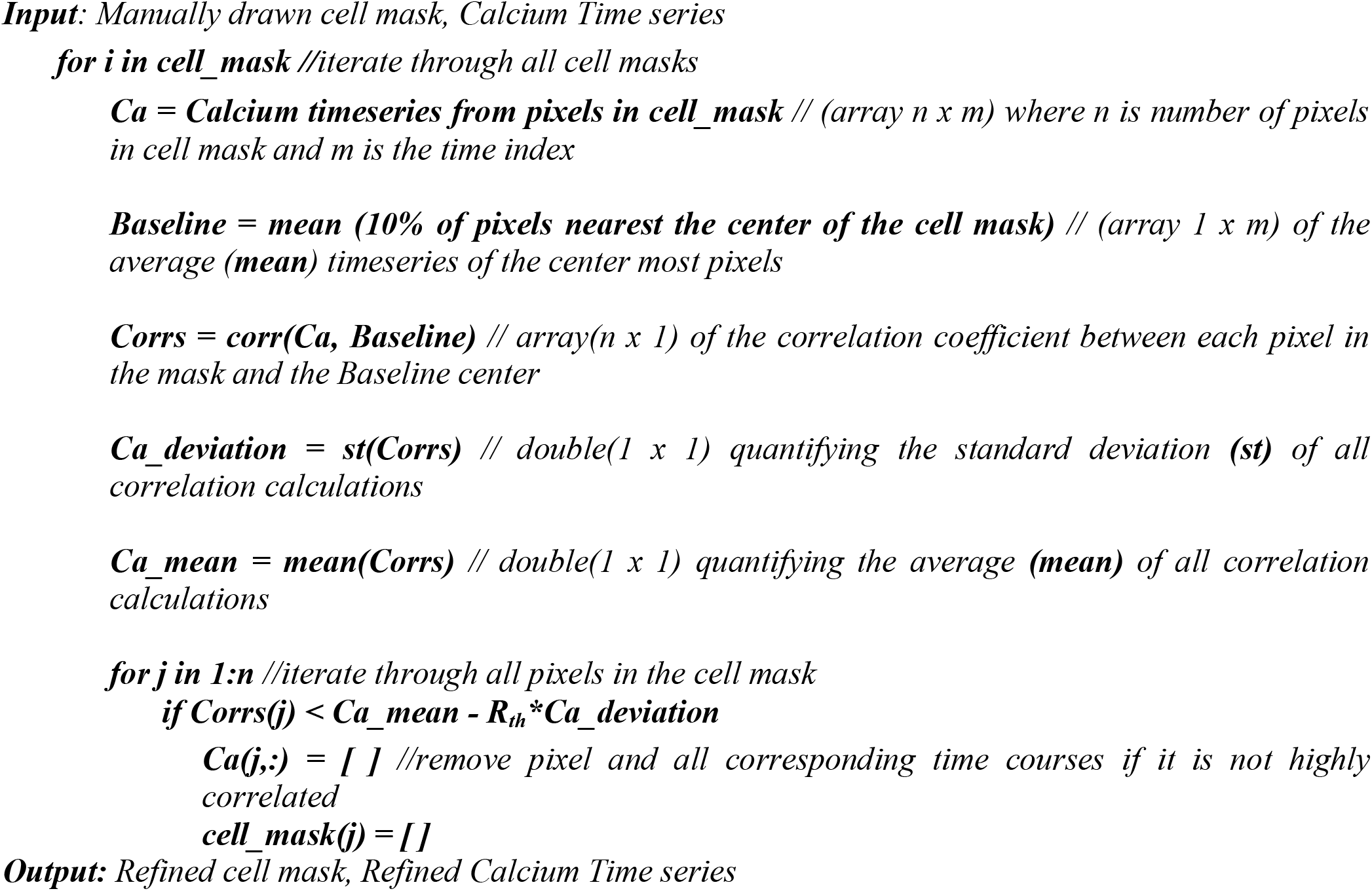

### Algorithm 2

*Algorithm for CRISP semi-minor axis identification*

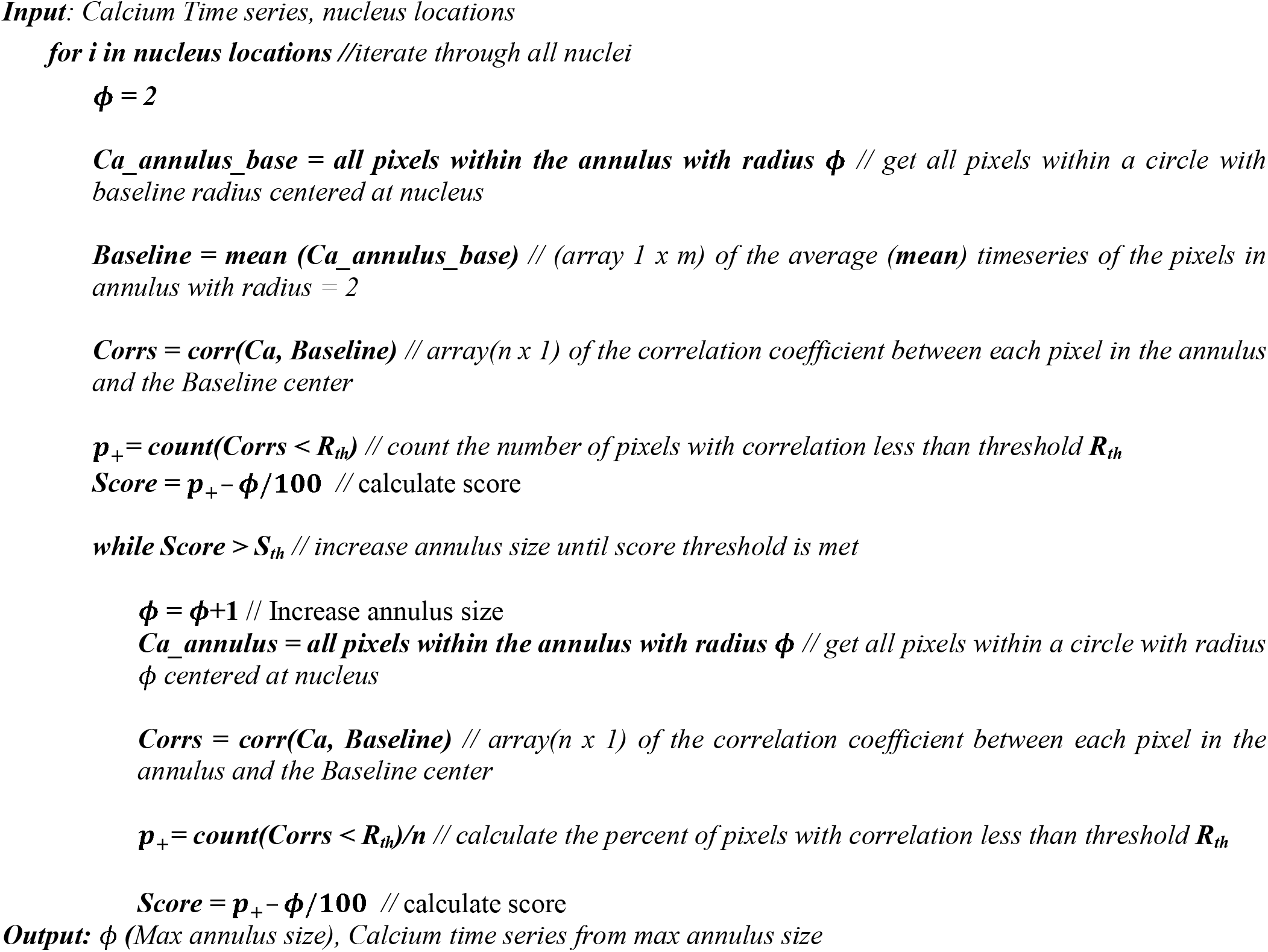

## Notes

### Competing Interest Statement

The authors have declared no competing interest.

### Summary of Updates

Updated abstract and figure 1 with additional information

